# Selection of time points for costly experiments: a comparison between human intuition and computer-aided experimental design

**DOI:** 10.1101/301796

**Authors:** Daphne Ezer, Joseph C. Keir

## Abstract

**Motivation:** The design of an experiment influences both what a researcher can measure, as well as how much confidence can be placed in the results. As such, it is vitally important that experimental design decisions do not systematically bias research outcomes. At the same time, making optimal design decisions can produce results leading to statistically stronger conclusions. Deciding *where* and *when* to sample are among the most critical aspects of many experimental designs; for example, we might have to choose the time points at which to measure some quantity in a time series experiment. Choosing times which are too far apart could result in missing short bursts of activity. On the other hand, there may be time points which provide very little information regarding the overall behaviour of the quantity in question.

**Results:** In this study, we design a survey to analyse how biologists use previous research outcomes to inform their decisions about which time points to sample in subsequent experiments. We then determine how the choice of time points affects the type of perturbations in gene expression that can be observed. Finally, we present our main result: NITPicker, a computational strategy for selecting optimal time points (or spatial points along a single axis), that eliminates some of the biases caused by human decision-making while maximising information about the shape of the underlying curves, utilising ideas from the field of functional data analysis.

**Availability:** NITPicker is available on GIThub (https://github.com/ezer/NITPicker).

## 1 Introduction

In many areas of experimental science, scientists are interested in the behaviour of some system under a wide range of conditions. For instance, a plant biologist might be interested in measuring gene expression in a set of mutant plant varieties under various environmental conditions, such as varying temperature, watering treatments and light intensities, in what is called a factorial (or multi-factor) experimental design (Figure 1A), but these types of experiments can be very expensive [Lundstedt *et al.*, 1998].

**Fig. 1.**
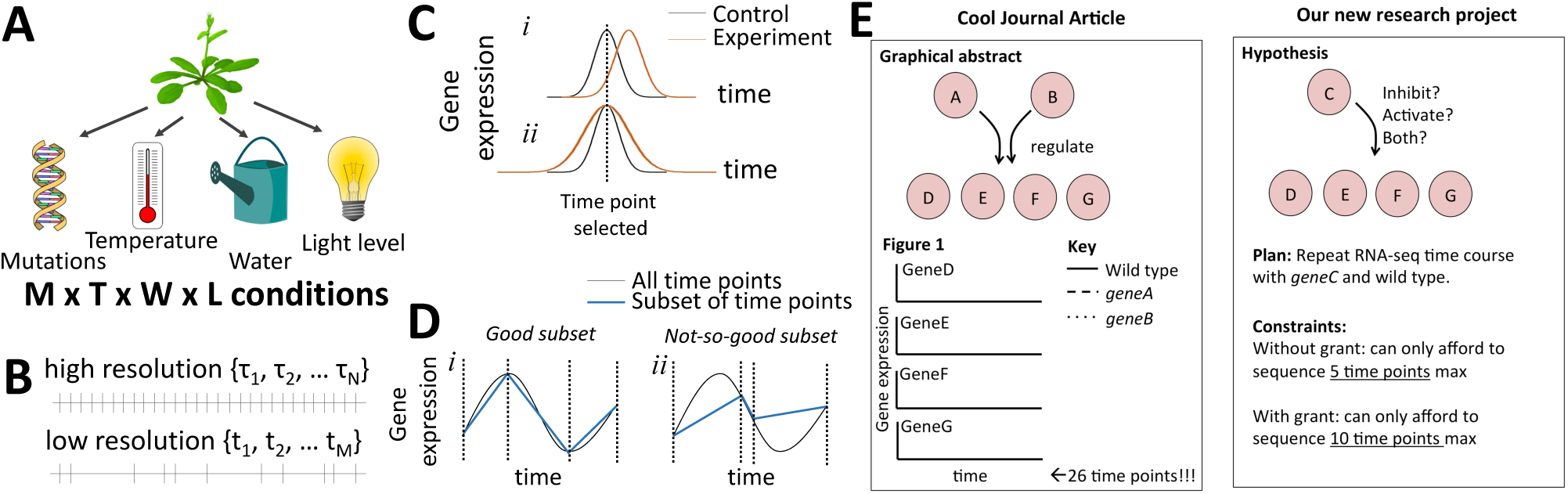
Time point selection is an important part of experimental design. (A) Here is an example of a multi-factor experimental design (B) Given a set of high resolution time courses, sampled at *τ_n_* we try to find a subset of time points **t**_*m*_ for future follow-up investigations. (C) The selection of time points will determine how a researcher will interpret a perturbation in gene expression. (D) In this paper, we define good time points as those that enable us to infer the shape of the function. (E) Biologists were presented with this figure to help explain their experimental design task.

To complicate matters even more, there are a large number of experiments which do not simply measure some discrete quantity, but instead aim to measure a *function*. Typically, researchers are interested in the behaviour of a quantity *over time* (or, in some cases, space). For example, many genes’ expression levels vary over time in intricate ways, especially since genes are often expressed in bursts. The shape of the burst provides insight into the regulatory mechanisms governing it [Nicolas *et al.*, 2017, Ezer *et al.*, 2016]. Even the degree to which a gene is sensitive to an environmental condition is often time-dependent; for instance, there are a different set of *Arabidopsis thaliana* genes that are sensitive to light at night and during the day [Rugnone *et al.*, 2013].

Ideally, a scientist would want to sample at a large number of time points under each experimental condition, but this might be infeasible, especially if the experiments are expensive to run. In such circumstances, the scientist might conduct a small number of high resolution time course experiments, and then use the information gathered to select a subset of time points for further investigation under the entire range of experimental conditions (Figure 1B). For example, many high resolution time course experiments have recently been published as part of large projects or consortia, including the high resolution time courses of fruit fly [Graveley *et al.*, 2011], roundworm [Boeck *et al.*, 2016], or mammalian lung development [Ardini-Poleske *et al.*, 2017], but a small lab that is interested in repeating the experiment under slightly different conditions might not be able to afford to use as many time points. Choosing the right subset of time points is clearly important from the point of view of accuracy, but it will also determine what *kinds* of gene expression perturbations can be observed in the follow-up experiments. For example, if a gene were to have a burst of expression in the first experiment, it would be reasonable for the biologist to select the time point corresponding to the peak of the burst for follow-up experiments (Figure 1C). Then, if the gene shifts the timing of its peak gene expression in the experimental condition, this change would be detected by the experiment, although it would not be possible to relate this to a change in the timing of the peak expression using this measurement alone. On the other hand, if the peak gene expression is the same, but the shape of the distribution changes, the biologist would not be able to detect any change at all (Figure 1Cii). Clearly, it would be beneficial to select time points that help us accurately determine the full gene expression profile (Figure 1D), while remaining sensitive to the expected types of perturbations. There have been many previous attempts to implement non-uniform sampling of time points using human intuition or heuristic-based strategies [Cortijo *et al*., 2017, Schulz *et al*., 2013], but as far as we are aware, there has been no systematic study of how biologists usually select time points for conducting their experiments and how these heuristics might affect their experimental outcomes.

In this paper, we ask 50 biologists to use prior information from simulated high resolution time course experiments to inform the selection of time points in follow-up experiments, and we analyse how their choices might affect the conclusions that can be drawn from the experiment (Figure 1E). Then, we develop a new statistical tool, called NITPicker, which selects informative time points for follow-up experiments given a set of example curves from a high resolution time course. NITPicker uses methods from functional data analysis to find these optimal points, and improves on current approaches for selecting time points for follow-up experiments. The growing field of functional data analysis is focused on developing new statistical techniques to analyse data sampled from continuous curves [Ramsay and Silverman, 2005, Wang *et al.*, 2015], In our case, in order to determine the relative importance of each time point for follow-up experiments, we need to know what types of curves we might observe under different experimental conditions. If all possible curves are equally likely to be observed in the new experimental conditions, then any set of time points would be equally sensible to select for the followup experiments. In reality, we can expect the observed curves to arise from some non-uniform probability density function of curves, whose parameters we must attempt to infer from the example curves that are available.

Some previous methods for selecting time points for follow-up experiments imagine that all the biological material is collected at each of the original time points and stored, but that the material from each time point is sequenced sequentially based on previous outcomes [Rosa *et al.*, 2012, Singh *et al*., 2005], However, this is rarely a practical experimental strategy, as it can result in a large amount of wasted time and effort, since biological material is collected at every time point, including those points which are not used in the later analysis. Also, sequencing in parallel can be much quicker and less expensive than sequencing sequentially.

The recent Time Point Selection (TPS) method developed by Kleyman *et al* [2017] is a substantial improvement in that it does not depend on this sequential experimental design strategy, and it considers the full shape of the gene expression profile, a strategy also used by NITPicker (Figure 1D). However, it has three downsides that might limit its use in practice. First, it uses a greedy search strategy for finding time points, which might be prone to finding local optima rather than global optima. In NITPicker, we identify that an optimisation problem described by Kleyman *et al* [2017] is in fact the same as a simpler problem in computer science – finding the shortest path through a directed acyclic graph – which can be solved directly by a dynamic programming algorithm (specifically, a modified Viterbi algorithm).

Second, TPS attempts to find the time points which lead to the most accurate fit *for the data in the training set*, so it might not generalise to new gene expression profiles that differ even slightly from the training set. As such, it is useful for follow-up experiments which attempt to repeat the original experiment at a lower resolution. However, in many experimental situations, we interested in selecting the time points which provide the most information about *how the curve changes* in experimental conditions. For this reason, in NITPicker we use the training set to develop a probability distribution of gene expression curves [Tucker *et al*., 2013], which allows us to address the slightly different (and more frequently encountered) question of finding the optimal time points for detecting and modelling *perturbations* of the data.

Third, TPS directly uses gene expression profiles from the high resolution time course to select the new time points, a strategy which is potentially vulnerable to experimental noise. On the one hand, we can imagine a scenario in which, for some period of time, the data is very noisy, before later settling down. In this case, TPS is almost certain to select time points in the noisy region (allowing us to more accurately model the noise), despite this providing very little useful information. On the other hand, individual anomalies in the data might cause TPS to select the associated time points. Since NITPicker uses a probability density over gene expression curves [Tucker *et al*., 2013], this decreases the risk of overfitting the training set, avoiding the latter problem. We also adapt NITPicker for use in the former scenario, by fitting the *inverse coefficient of variation* rather than the data itself. In summary, NITPicker does not get trapped in local optima, addresses a wider range of experimental design questions, and is less sensitive to noise in the training set.

This paper demonstrates that human intuition can lead to sub-optimal experimental design decisions, but through computer-aided experimental design strategies, it is possible to make more informed experimental choices. In particular, we present a tool which allows researchers to select optimal time points for follow-up experiments in a range of different experimental design situations.

## 2 Results

### 2.1 Experimental design survey

We developed a survey to help us determine how biologists use high-throughput experimental data to inform their time point selection in followup experiments (Figure 1E, S1). Fifty biologists from a variety of career stages and expertise (Figure 2A-C) were presented with a series of four experimental design scenarios. In each case, they were presented with simulated curves displaying gene expression over time for four genes, in three experimental conditions, sampled at 26 time points. After viewing these figures (Figure 1E, S1), biologists were asked to select a subset of time points at which they would measure the gene expression of the same genes in different experimental conditions. For each example, the biologists were first asked to select 5 time points, then a set of ten time points that they would select ‘in the case they win a grant’.

**Fig. 2.**
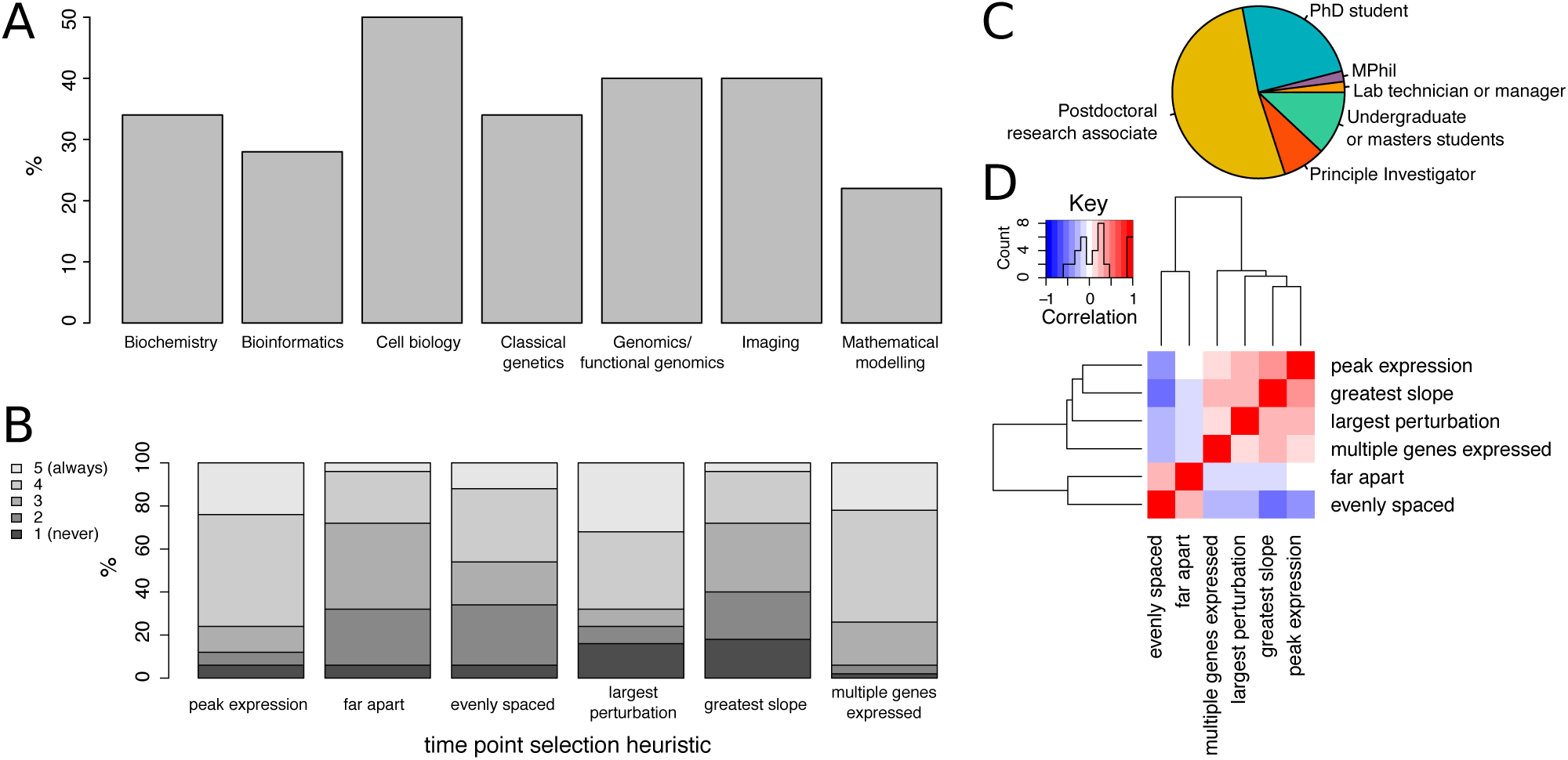
Here we present survey demographics. (A) Survey participants indicated familiarity in a variety of sub-disciplines of biology. (B) They rated heuristics they used to select the time points, from 1 (never) to 5 (always). These statements were phrased as: ‘I preferred to select time points …(i) with maximal gene expression, (ii) that were far apart from one another, (iii) that were evenly spaced, (iv) that had the largest gene expression differences between the samples, (v) in areas where the slope of the gene expression curves were steepest, (vi) where more than one gene was expressed. (C) Participants came from a variety of career stages. (D) Hierarchical clustering of the correlation coefficient between each pair of heuristics in B.

The simulated examples presented to the biologists were all drawn from the skewed Gaussian function – see Table S1. We chose skewed Gaussian distributions as they are a good model for the bursts of expression previously observed in biological systems of interest, including G-box regulated genes expressed at dawn [Ezer *et al.*, 2017], and temperature responsive genes [Cortijo *et al.*, 2017]. In each of the four scenarios presented to the biologists, one of the parameters defining the skewed Gaussian was changed, while the others were kept the same. More specifically, the scenarios equate to changing the mean, the standard deviation, the skew, and the amplitude of the gene expression curves.

Afterwards, biologists answered questions about the heuristics they deployed when selecting time points (Figure 2B). Three of the most popular self-described strategies for selecting points were selecting areas of peak gene expression, selecting points where there was the largest perturbation and selecting points where multiple genes were expressed. Those who said that they chose points that were evenly distributed were more likely to say that they choose points that were far apart from one another (Figure 2D). Overall, these survey results suggest that there are many diverse strategies that biologists use to select subsampled time points for followup experiments, and there is no consensus regarding which strategy is best.

### 2.2 Defining good time points

In order to compare the strengths and weaknesses of each heuristic, we need to make a clearer definition of what constitutes a *good* time point to select. One possible strategy would be to try to select a set of time points that are best able to distinguish the shape of the curves. For this we select a criteria very similar to that presented by Kleyman *et al.* [2017] in that we want to minimise the *L*^2^-distance between the curve generated by sampling at all the time points, and the curve generated by sampling only at a subset of time points. However, Kleyman minimises this distance over all curves in the training set – instead, we use the training set to generate a probability density over the space of curves, and then minimise the expected distance. Some of the advantages of this approach have already been mentioned in the introduction.

Suppose that we have an initial high resolution time course, with data sampled at times ***τ*** = {*τ*_1_, *τ*_2_, …*τ_N_*}. We must select a subset of these time points, which we will call ***t*** = {*t*_1_, *t*_2_, … *t_M_*}, so as to minimise the expected error:

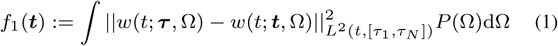

where *w*(*t*; ***t***, Ω) is a gene expression function evaluated at the time *t*, parameterised by a set of parameters Ω and interpolated between time points in the set ***t*** (either through a linear interpolation or spline), and *P*(Ω)dΩ is a probability measure on the space of parameters (i.e. *P*(Ω) is the probability density associated with the set of parameters Ω). We use the standard notation for *L*^2^ norms, that is, given a function of time *w*(*t*) we define

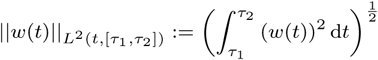

In many cases we are not necessarily interested in the shape of the curve, but rather the difference between the control and an experimental condition. Let *g*(*t*, Λ) be the gene expression curve in the control condition, parameterised by Λ and sampled at time points *t*. Then we want to minimize the expected error in the difference:

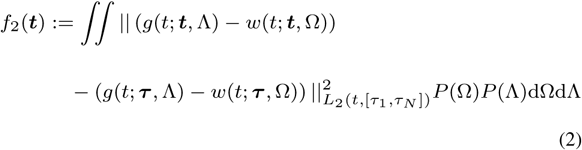

If there is only one ‘control’ curve (for instance, if the scientists have not included replicates), then there would only be one possible value for the parameters Λ and the equation is simplified to:

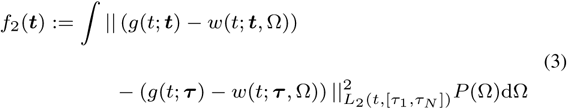

If there are periods of time with different amounts of random variability, then we might wish to sample less frequently in areas that have lots of variability – we might accept having less accuracy in predicting the shape of the curves in noisy regions if we can accurately model their shapes in regions with less noise. To accomplish this, we should attempt to find the shape of the curve representing the difference between the control and experimental conditions *normalised by the variance.* In other words, we minimise the expected error in the *inverse of the coefficient of variation:*

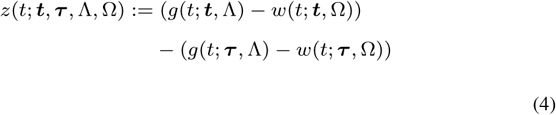

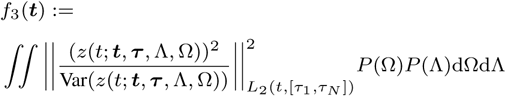

where the variance is itself a function of *t*: it is the variance of the function *z*(*t*; ***τ***, ***t***, Λ, Ω) with respect to the probability measure *P*(Ω)d*P*(Λ)dΛ.

Within the experimental design survey, the gene expression curves were drawn from a known probability distribution (i.e. the curves were skewed Gaussian functions, with the parameters either fixed or drawn from a Gaussian distribution), so we can quantitatively compare the time points suggested by the survey participants by directly inputting each set ***t*** of time points suggested by a survey participant into the above functions. Based on the wording of experimental design scenario as presented in the survey (with a clear distinction between a control and experimental condition), minimising the function *f*_2_ probably corresponds most accurately with the goals of the biologists in the follow-up experiment.

Of course, there are many other criteria that might be used to determine the “optimal” set of time points to select, but for the purpose of this manuscript we focus on these three criteria, as they are intuitive, relatively simple, and – as we shall see later – the best solution can be computed exactly with a dynamic programming algorithm. Note also that our algorithm can easily be adapted to deal with many other criteria. For example, if it is important to accurately measure both the curve *and its first s* derivatives, then we can simply replace the *L*^2^ norms in the above expressions with the norms associated with the Sobolev spaces *H^s^*.

#### Factors associated with good time point selection among biologists

We can now evaluate how the choice of heuristics used by biologists correlates with their *f*_2_ score. First, for each experimental design scenario, we calculated the difference between the control curve shown in the survey and 1000 new gene expression curves sampled from the same distribution of curves used to generate the survey questions. Next, we calculated the *L*^2^-distance between the curve generated by sampling at all time points and the curve produced by sampling only at the suggested subset of time points. Since there were four experimental design scenarios presented in the survey, each with four genes, we had a total of sixteen *L*^2^-distances for each survey participant. In order to provide equal weight to each of these sixteen conditions, we ranked the participants within each of the 16 conditions, and then calculated the average rank.

The participants’ average rank when selecting 5 time points was correlated with their rank when selecting 10 time points (Figure 3A). Next, we used regularised linear regression to determine whether the heuristics for selecting time points were correlated with the average rank of the participants. The λ parameter was determined by cross-validation – the value of λ that minimised the mean-squared error was chosen. We found that selecting points that are far apart, evenly spaced, or points that overlap between multiple gene expression curves were all heuristics that improved the average rank. On the other hand, the average rank was negatively affected by selecting time points where there was the largest different between the the conditions (Figure 3B-C). Qualitatively, it appears that biologists tend to select time points near the centre of the distributions, rather than the beginning or end of the time course (Figure S2). This heuristic might have a negative effect on the biologists’ scores, since in order to have an accurate idea about the shape of the distribution it is important to have an idea of what the curves are doing at either end of the time course. It may be that biologists are assuming that the “tails” of the gene expression have a standard shape which is known *a priori*, and so sampling at these time points does not provide much information.

**Fig. 3.**
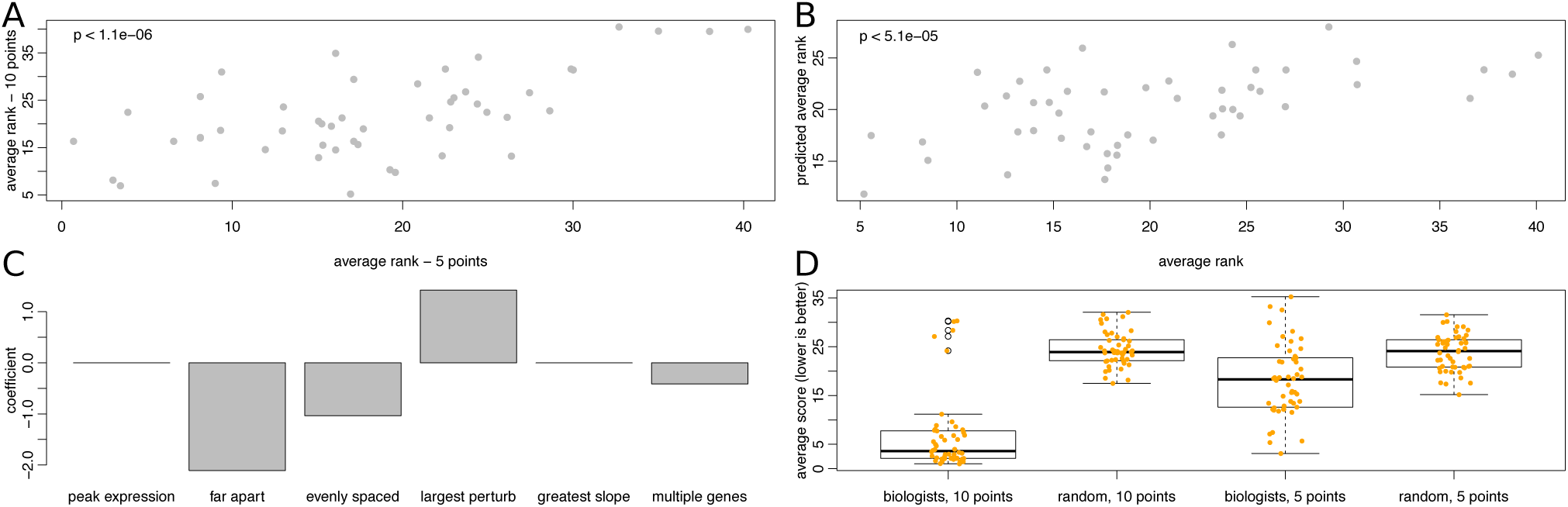
Results of the experimental design survey (A) For each experimental design scenario, biologists were asked to select a set of 5 time points and then a set of 10 time points for follow-up experiments, and we compare the average rank of each biologist in these two cases. (B) Regularised linear regression was used to determine how the heuristics in Figure 2B correlated with the average rank (combining the 5 time point and 10 time point average ranks). Here the actual average rank is compared to the predicted average rank. (C) The coefficient values in the model for (B) are shown here. Note that having a low rank indicates that the time points selected were good, so a negative coefficient indicates that heuristic is predictive of a good selection of time points. (D) Finally, we compare how the biologists performed relative to a random selection of points. We did this by making 50 random selections of time points, and ranking each biologist relative to this set of 50 time point choices. We also scored fifty new randomly selected time points, to serve as the null distribution, which we compared with a two sample, one-sided T-test. We found that biologists were significantly better than random, both when selecting 10 time points (*p* < 2.2*e* – 16) and 5 time points (*p* < 3.0*e* – 06).

While the biologists performed significantly better than random in both the 5-time point and 10-time point cases, but the improvement in score is much greater in the 10-time point case (Figure 3D), the topics of expertise did not seem to affect the outcome of the survey, although there is a slight anti-correlation between the level of seniority and the final score (i.e. undergraduates and masters students performed slightly better on average than principle investigators)– see Figure S3.

This analysis can provide us with some rules of thumb that might be useful for selecting time points, emphasising, for example, the importance of choosing time points that are far apart from one another or that have non-zero gene expression values for multiple genes. However, it is unclear whether these suggestions are useful *in general*, or whether they are specific to the type of experimental design scenarios presented as part of the survey. Instead, it would be best if it were possible to identify reasonable time points in a more consistent and systematic way. 70% of survey participants said that they would always or often prefer to use an algorithm for making these types of experimental design decisions, suggesting a real need for computational strategies for time point selection.

### 2.3 Defining a non-parametric probability density function of curves

In the experimental design survey, the probability distribution from which the gene expression curves were drawn was known explicitly, since we generated the gene expression curves ourselves. However, in real experiments this will not be the case. In order to effectively find the subset of time points that minimise *f*_1_, *f*_2_, or *f*_3_, we need a method to derive, from a set of example curves, the probability of observing particular curves in future experiments.

First, suppose that we have chosen some way of parameterising curves using a set of parameters Ω. Let *g_a_*(*t*) be the curve produced in the a-th high resolution time course experiment, and let Ω_*g_a_*_ be the associated parameters. Given the whole set of such parameters associated with all the high resolution time courses, we want to generate a probability density function *P*(Ω) on the parameter space.

In some cases, a scientist might have a model in mind that describes the functions – in this case they can directly fit the parameters. However, in most cases we’ve encountered scientists do not have such a model, nor can one be justified on theoretical grounds, so we need a *non*-*parametric* way of defining a probability distribution of functions given a set of examples. In the discipline of functional data analysis, techniques have been developed to define these probability distributions – see Tucker *et al.* [2013] – a process that involves first aligning the functions (“registration”) and then parameterising the horizontal and vertical shifts (using functional Principle Component Analysis). For completeness, we will summarise their protocol below.

First, we take a set of known functions (our set of high resolution functions, *g_a_*(*t*)) and *align* them. In order to effectively align the gene expression curves in a shape preserving way, we define a distance between curves in terms of the square root slope function (SRSF):

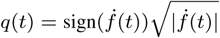

Note that, given *q*(*t*) and the initial value *f*(0), we can recover the corresponding function *f*(*t*) via
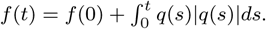

Now, we define the *y*-distance between *f*_1_ and *f*_2_ as

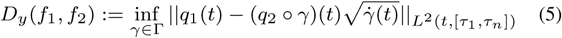

where *γ* is a function defining the amount of *x*-axis warp, and Γ ⊂ *L*^2^ is the set of *warping functions*. *γ* ∈ Γ must have some special properties: (i) *γ*(*t*) ∈ [0, 1] (ii) *γ* is monotonically increasing (i.e its slope is positive) (iii) ∥*g*(*t*)∥_*L*^2^(*t*)_ = 1. Because of this last point, the warping functions must lie on the *unit sphere in L*^2^, which can be thought of as an infinite-dimensional sphere.

Given a function *g*_(mean)_ (see Tucker *et al.* [2013] for the appropriate *g*_(mean)_), a dynamic programming algorithm is used to find a vector of functions ***γ*** = (*γ_a_*(*t*)), where *γ_a_* corresponds to the warping function found when computing *D_y_* (*g*_(mean)_, *g_a_*) (see equation 5). It also provides us with a set of aligned functions *f_a_* and a corresponding set of aligned *q* functions *q_a_*.

We would like use the set of warping functions *γ_a_* to define a probability density function on (some subset of) Γ. However, Γ is not a linear space: given two warping function *γ*_1_ and *γ*_2_, their sum *γ*_1_ + *γ*_1_ cannot be interpreted as a warping function, since it will not lie on the unit sphere in *L*^2^. Hence, we cannot immediately apply functional Principle Component Analysis. Instead, we linearise the space Γ by first finding the centroid of the points *γ_a_* on the surface of the sphere (the “Kercher mean”, *γ*_(mean)_). Note that this is itself a function of *t*. Next, we use the *exponential map* at *γ*_(mean)_, which provides us with a map from the tangent space at *γ*_(mean)_ (which *is* a linear space) to the sphere itself. More precisely, given a tangent vector to the sphere, we find the point on the sphere reached by exponentiating this tangent vector, using the Lie-group structure (under composition) of the unit sphere in *L*^2^. In this way, we can associate a vector in the tangent space at *γ*_(mean)_ to each of the warping functions *γ_a_*. Now, to decrease the dimensionality of the space (and to decrease the number of free parameters in the model), we can perform a functional Principle Component Analysis (fPCA) of this linearised space, then fit an independent normal distribution along each principle axis. This gives us a probability density on the tangent space. Finally, we can use the exponential map again to map this probability density function directly onto the sphere.

An fPCA can also be performed on the the y-axis deformations. This time, the functions defining the *y*-axis deformations are simply functions in *L*^2^, which is already a linear space, so we don’t need to perform the linearisation step.

In summary, this strategy results in Ω being defined as the space of fPCA coordinates, associated with both *x*- and *y*-deformations. *P*(Ω) is then given by the multivariate normal distribution with a diagonalised covariance matrix.

To estimate the integrals involved in the definitions of *f*_1_, *f*_2_ or *f*_3_, we take *J* samples from the probability distributions defined above. For example, we estimate *f*(***t***) as

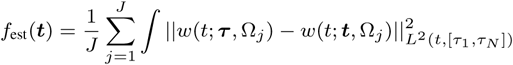

where Ω_*j*_ is the set of parameters corresponding to the *j*-th sample drawn from *P*(Ω).

The benefit of this approach is that it allows the user define very complicated functions for *w*(*g*, Ω) and *P*(Ω), and still be able to apply NITPicker. The downside of this approach is its random nature, which means that we don’t know the error between the actual value of the integral and this estimate, although we can be confident that *f*_(est)_ is close to *f* if *J* is sufficiently large.

### 2.4 NITPicker algorithm

A dynamic programming algorithm can be employed to find the set of *m* time points ***t*** that minimise *f*(***t***) (where *f* = *f*_1_, *f*_2_ or *f*_3_). In essence, the problem is identical to finding the path that minimises the distance in a directed acyclic graph that contains exactly *m* edges, which can be calculated with a modified Viterbi algorithm (Figure 4A). Consider a graph with *N* + 2 ordered nodes – a ‘start’ node, *N* nodes that represent each time point in the high resolution time course, and an ‘end’ node. For ease of notation, we index the start node with 0 and the end node with *N* + 1. Each node is connected by edges that point to all the nodes that are ahead of it, and we set the value of the edge joining node *i* to node *k* to be:

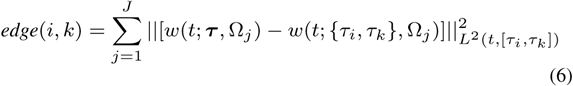

**Fig. 4.**
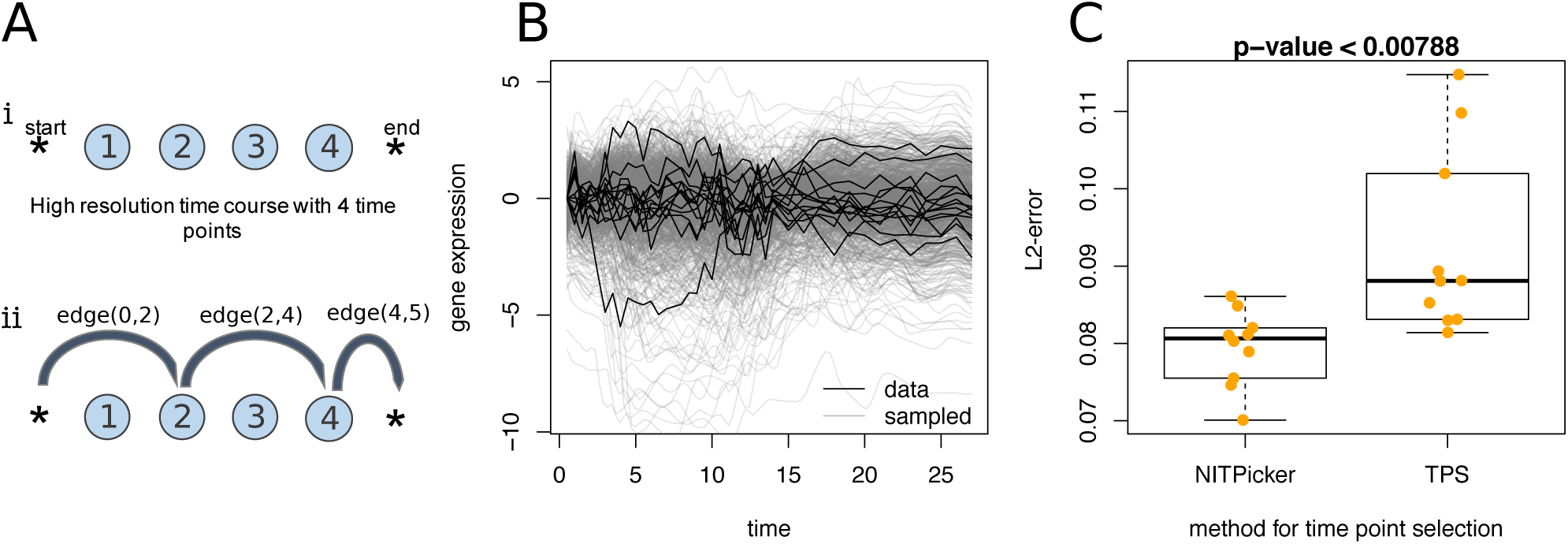
(A) The time points in a high resolution time course are represented by nodes. In the case shown (i), there are four time points in the high resolution time course. (ii) In the follow-up experiment, time points 2 and 4 are chosen, corresponding to a particular path through the network. The length of this path is the sum of edge(0, 2), edge(2,4) and edge(4,5). (B) The lung dataset was collected in Kleyman et al. [2017] and used to test their TPS algorithm. We split this data into ten groups, with each group consisting of 14 or 15 curves. One such group is shown in black. This is then used to train NITPicker: the grey curves show a population of curves generated by NITPicker modelled on the data shown in black. (C) The same set of 14 or 15 curves is used to train TPS and NITPicker, which each select 8 timepoints (as in the example of Kleyman et al. [2017]). We then test this selection by calculating the *L*^2^-error encountered in the remaining 9/10ths of the dataset. We repeat this for each of the ten groups. The errors found using NITPicker’s choice of time points are consistently lower than the errors produced using the timepoints selected by TPS.

In other words, the value of the edge joining node i to node *k* is the *L*^2^-error caused by selecting times *τ_i_* and *τ_k_* and none of the times in between.

Now we need to find the shortest path with *K* edges that goes from the start node to the end node. An *N* by *N* by *K* table can be assembled where each element is:

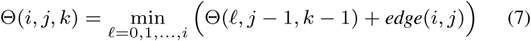

and Θ(0, *j*, 0) = 0, Θ(*i* > 0, *j*, 0) = ∞. In other words, the value of Θ(*i*, *j*, *k*) is the minimum error when going from *τ*_0_ to *τ_i_* and then immediately to *τ_j_*, using exactly *k* edges.

The value of *f*_est_(***t***) is Θ(*N* + 1, *N* + 1, *K*). As we construct the table Θ, we also save another matrix Θ_*min*_, with entries given by Θ_*min*_(*j*, *k*) = min_*i*_ Θ(*i*, *j*, *k*). The value of Θ_*min*_(*j*, *k*) is then the minimum *L*^2^-error when going from *τ*_0_ to *τ_j_* using *k* steps. We also construct a similar *N* by *K* matrix Θ_*trace*_ (*j*, *k*), with entries given by the value of *i* which minimises Θ(*i*, *j*, *k*). This can then be used to find the time points: we set *t_k_* = Θ_*truce*_(*N* + 1, *k*), and then *t*_*k*–1_ = Θ_*trace*_(*t_k_*,*k* – 1).

Note that the value of *edge*(*i*, *j*) actually depends on the previous *R* time points if a spline of degree *R* is used. However, the index of the previous *R* best nodes given edge (*i*, *j*) can be easily computed from the traceback matrix, although this makes NITPicker run much more slowly. Furthermore, using a spline of degree greater than one can produce edge-effects, especially in the beginning of the sequence as we use the deBoor algorithm to calculate the spline [R. and de Boor, 1980]. This problem can be reduced by running the dynamic programming algorithm twice – once forward and once backwards.

### 2.5 Testing NITPicker on real world data

In order to determine how well NITPicker works in practice, we apply it to three real-world datasets, and address a slightly different experimental design question in each case, corresponding to *f*_1_, *f*_2_ and *f*_3_. First, we test the performance of NITPicker when minimising *f*_1_, which means that we are trying to accurately model the shape of the curves. As this is the same problem proposed by Kleyman *et al.* [2017], we compared the performance of NITPicker to their TPS method, using the same lung gene expression dataset. In this case, we are presented with an RNA-seq experiment in only one experimental condition, but containing a large number of genes. The genes were hand-selected by Kleyman et *al.* [2017] as they are believed to be involved in lung development. As such, a reasonable hypothesis appears to be that the corresponding gene expression curves are all drawn from a single probability distribution over curves. To determine if this is an accurate model, we randomly split the dataset into 10 roughly equal partitions, each containing 14 or 15 gene expression curves. We then use these 14 or 15 curves to select eight time points using either NITPicker or TPS, and then evaluate how well we can model the remaining nine-tenths of the gene expression curves by sampling only at these eight points. This evaluation is carried out by calculating the *L*^2^ -error that arises from only sampling at the eight time points selected, rather than at every point. We observe that NITPicker comes up with a sensible probability distribution of gene expression curves using only 14 curves (Figure 4B) and performs significantly better than TPS (Figure 4C). Note that TPS always selects the first and last time point, so it has less flexibility in selecting time points, and this might account for some of the difference in *L*^2^-error.

In functional data analysis, two of the standard datasets for testing new algorithms are the Canada weather dataset [Ramsay and Silverman, 2005] and the Berkeley growth dataset [Tuddenham R.D., 1954], both of which provide additional examples of real world data with a functional form.

The Canada dataset contains the average temperature measured each month across a number of Canadian cities, and we use this to test NITPicker’s performance when minimising *f*_2_. Suppose that we are interested in the difference in weather between cities in Canada and Resolute, one of its most northerly and coldest cities. In other words, we will be finding the value of *f*_2_, where *g* is the weather of Resolute and *w* is the weather of the other Canadian cities. Suppose (for the sake of illustrating our methods) that we are in a position to sample the weather patterns in some new cities in Canada, but we can only afford to measure the weather in 5 of the 12 months. The raw data is shown in Figure 5A, showing that we can generate a sensible probability density function for temperature curves in Canada. The best months to sample according to NITPicker are drawn as vertical lines.

**Fig. 5.**
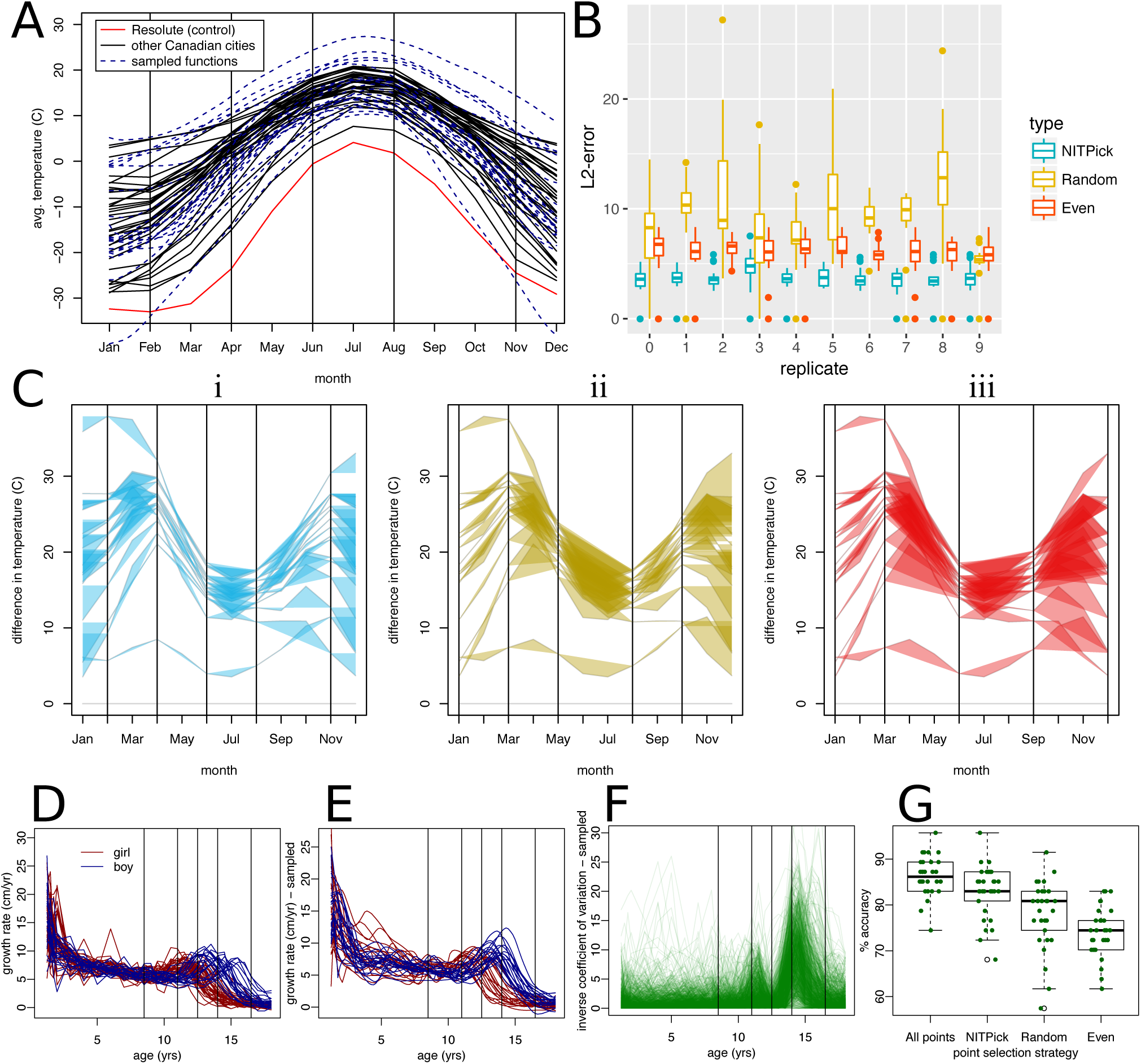
Applying NITPicker. (A) Monthly temperatures from a set of cities in Canada are shown in black. For the purpose of this paper, we consider the ‘control’ condition to be the temperature of Resolute, Canada, which is shown in red. A probability distribution of curves was constructed on the basis of the temperature curves for all cities – curves sampled from this probability distribution are indicated in dashed-blue lines. The vertical lines represent the ‘best time points’ to sample from, according to NITPicker. (B) For each of ten replicates, we selected the time points to sample using half the city curves, and scored the selection of time points on the other half of the city curves. For each city in the test set, we calculated the *L*^2^-error between the curve generated by sampling every month and the curve generated by linear interpolation between the selected subset of time points. (C) Here we present an example of how we evaluate a test set for a selection of time points (vertical bars) selected by NITPicker (i), random (ii), and evenly sampled (iii). The coloured-in area displays the error that arrises from sampling only at the designated time points. (D) The growth rate of boys and girls from the Berkeley growth dataset were used to develop probability distributions of curves for boys and girls, with sampled curves shown in (E). (F) We were interested in estimating the shape of the inverse coefficient of variation, shown in this figure. The selected time points are shown as vertical bars in D-F. (G) We used half the boy curves and half the girl curves to select time points to sample from, and to train a DD-classifier [Cuesta-Albertos et al., 2016,Li et al., 2012], and then calculated the percent accuracy on the other half of the boy and girl curves. This procedure was repeated 30 times with each method of selecting time points (selecting all the time points, 5 time points with NITPicker, 5 time points randomly, and 5 time points evenly).

To test the accuracy of the approach, we randomly split the cities into two equally sized groups – a training set and a testing set. We use the training set to select a subset of time points using NITPicker, and then we test the strength of these time points on the test set of cities. More specifically, we evaluate the *L*^2^-distance between the curves sampled at all points and those sampled only at the time points selected by NITPicker. We find that NITPicker-selected time points perform better on the testing set than either randomly sampled points or evenly sampled points (Figure 5B), an example illustrating this is shown in Figure 5C. This demonstrates that NITPicker can be used successfully to select time points that help distinguish between a control curve and a distribution of curves from experimental conditions.

The third dataset represents growth data from a group of boys and girls (Figure 5D) – despite the unusual shape of the curves, it is possible to develop a reasonable probability distribution of growth curves (Figure 5E). The largest variance in growth rates is found in the early years; however, from the point of view of distinguishing the two populations, the most informative difference in growth rates between boys and girls is seen during adolescence. Suppose that we want to sample at time points that can help us accurately determine the difference in growth rates between girls and boys. In other words, we don’t mind if the shape of the curve is less accurate in periods of time with lots of variability, but we wish to accurately estimate the shape of the difference between girls and boys in periods of time with less variability within each population – for this we are interested in minimising *f*_3_ (see Figure 5F). The point of this exercise is to select time points that help us estimate the shape of the difference between girl and boy curves; however, as by-product of the procedure we might hope that we can select time points that are reasonable at predicting whether an individual growth curve comes from a boy or a girl. Similar to our analysis for the Canada dataset, we split the curves into training and testing sets, but this time we not only select a set of time points using the training set, but we also train a classifier commonly used to classify functional data [Cuesta-Albertos *et al.*, 2016, Li *et al.*, 2012]. Although, as expected, the best classifier used all the time points, NITPicker-selected time points could be used to develop a more accurate classifier than selecting time points either evenly or randomly (Figure 5G).

### 2.6 Comparison between human intuition and computer-aided experimental design

Finally, we can compare the time points that were selected by NITPicker to those selected by human biologists (Figure 6A-B, S3). Even though there are only three examples of each gene, we found that we can estimate sensible probability density functions of curves for each example (Figure 6A). The time points selected most often by humans are also often selected by NITPicker (i.e. see time points 10, 15, and 17 in Figure 6B). The full results when selecting 5 time points are shown in Figure 6C, and the results for 10 time points are in Figure S4. NITPicker performed better than most human biologists (its average rank is 18.25 out of 51 when selecting 5 time points, and 13.56 when selecting 10 time points) – see Figure S5. While NITPicker does better than most humans, there are some biologists who consistently performed better than it, although there are not enough experimental design scenarios in the survey to determine whether this is by luck, or whether some scientists are using intuitive time point selection strategies that are not captured by NITPicker.

**Fig. 6.**
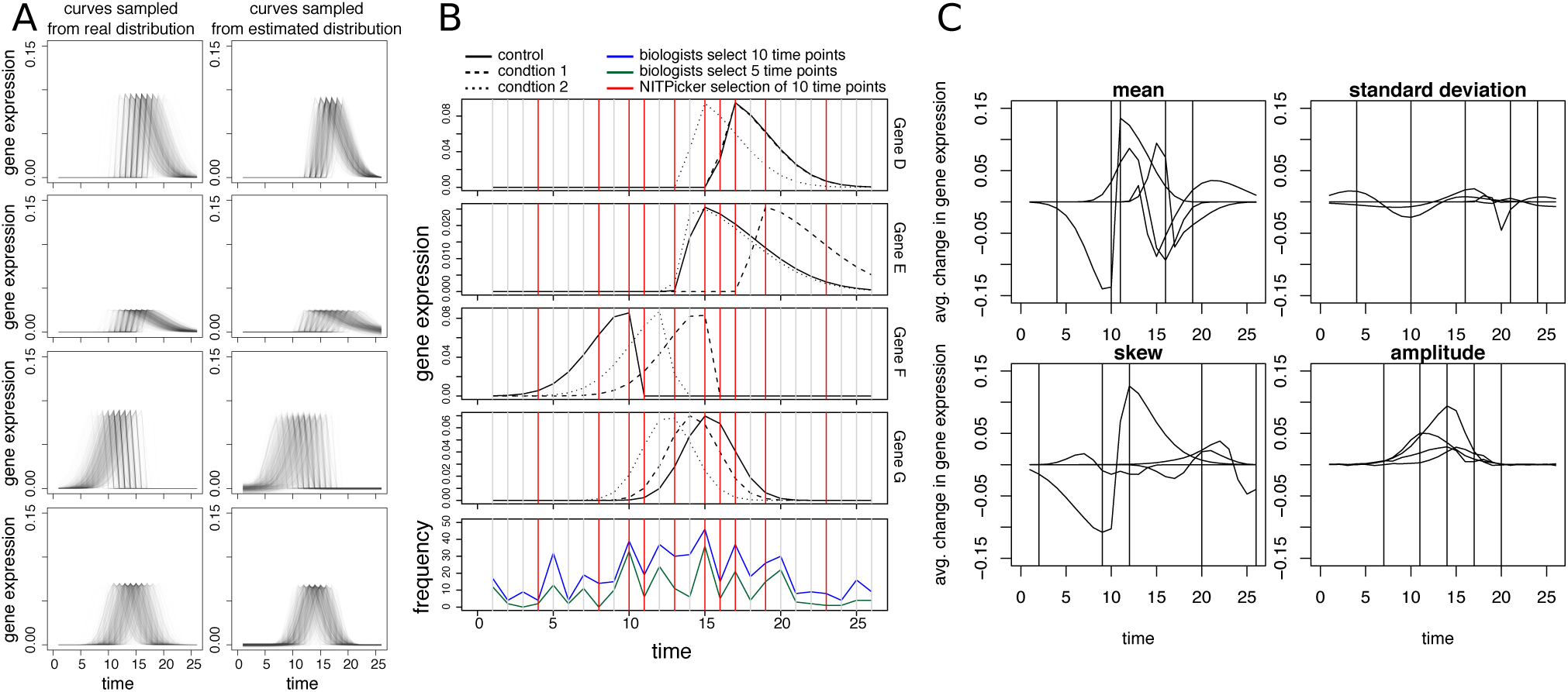
Testing NITPicker on the survey data. (A) The left column shows curves drawn from the populations of skewed Gaussian curves used in one of the experimental design scenarios in the survey. The right hand column shows populations of curves generated by NITPicker, using as input only the three examples of curves shown, for each gene, in (B). This figure represents the information provided both to survey participants and to NITPicker: for each gene, three curves were provided: a control, and two perturbations of the control, generated by sampling one of the parameters for the skewed Gaussian from a normal distribution (in this case, the parameter being varied is the mean). At the bottom of (B) we plot the frequency at which different time points were chosen by biologists. Throughout (B), the vertical red lines indicate the 10 time points selected by NITPicker. (C) We plot the average difference between the control curve and the perturbed curves, for each experimental design scenario in the survey (there are four genes per scenario). The 5 time points selected by NITPicker are displayed as vertical black lines.

## 3 Discussion

In this paper we have presented NITPicker, an algorithm for selecting a subset of time points in time course experiment in a variety of experimental design situations. In contrast to previous strategies [Kleyman *et al.*, 2017, Rosa *et al.*, 2012, Singh *et al.*, 2005], NITPicker takes full advantage of the *functional* nature of the data to produce a non-parametric probability distribution over curves [Tucker *et al.*, 2013], which is then used to select the optimal time points. This approach minimises the risk of over-fitting the data, while also being better adapted to the situation in which the new time points are being selected for use in experiments which are run under *different conditions* to those used to collect the original data. We also use a dynamic programming algorithm to select the optimal time points, which provides an efficient method that is guaranteed to find the optimal solution (unlike the greedy algorithm used in TPS [Kleyman *et al.*, 2017]). We tested NITPicker on a variety of simulated and real-life datasets, and demonstrated the flexibility of this tool by addressing different experimental design questions in each case.

Our survey of biologists indicated a desire for a computational tool to help in making these important, and frequently encountered experimental design decisions. NITPicker provides such a tool, which is both flexible, accurate and can help to avoid biassed results which can result from *ad hoc* human decisions.

## Acknowledgements

We would like to thank the survey participants at the Sainsbury Laboratory in University of Cambridge and the Cambridge plant science department, and Dr. Philip Wigge for helping encourage survey participation.

